# TreeTime: maximum likelihood phylodynamic analysis

**DOI:** 10.1101/153494

**Authors:** Pavel Sagulenko, Vadim Puller, Richard A. Neher

**Affiliations:** Max Planck Institute for Developmental Biology, 72076 Tübingen, Germany; Biozentrum, University of Basel, Switzerland

## Abstract

Mutations that accumulate in the genome of replicating biological organisms can be used to infer their evolutionary history. In the case of measurably evolving organisms genomes often reveal their detailed spatiotemporal spread. Such phylodynamic analyses are particularly useful to understand the epidemiology of rapidly evolving viral pathogens. The number of genome sequences available for different pathogens, however, has increased dramatically over the last couple of years and traditional methods for phylodynamic analysis scale poorly with growing data sets. Here, we present TreeTime, a Python based framework for phylodynamic analysis using an approximate Maximum Likelihood approach. TreeTime can estimate ancestral states, infer evolution models, reroot trees to maximize temporal signals, estimate molecular clock phylogenies and population size histories. The run time of TreeTime scales linearly with data set size.

---

Phylogenetics uses differences between homologous sequences to infer the history of the sample and learn about the evolutionary processes that gave rise to the observed diversity. In absence of recombination, this history is a tree along which sequences descend from ancestors with modification. In general, the reconstruction of phylogenetic trees is a computationally difficult problem but efficient heuristics often produce reliable reconstructions in polynomial time (Felsenstein, 2004; Price et al., 2010; Stamatakis, 2014). Such heuristics become indispensable for large data sets of hundreds or thousands of sequences.

In addition to tree building, many research questions require parameter inference and hypothesis testing (Drummond et al., 2012; Pond and Muse, 2005). Again, exact inference and testing with large data sets is computationally expensive since it requires high-dimensional optimization of complex likelihood functions or extensive sampling of the posterior. Efficient heuristics are needed to cope with the growing data sets available today.

Phylogenetic tree reconstruction relies on substitutions that accumulated between the common ancestor and the sequences in the sample. Zuckerkandl and Pauling (1965) hypothesized that changes in protein sequences accumulate linearly and that the number of differences between homologous sequences can be used as a *molecular clock* to date the divergence between sequences. The molecular clock has since been used to date the divergence of ancient proteins billions of years ago as well as the spread of RNA viruses on time scales less than a year (Langley and Fitch, 1974; Rambaut, 2000; Sanderson, 2003; Yoder and Yang, 2000). Beyond dating of individual divergence events of the time of the common ancestor, algorithms have been developed to infer a tree where branch lengths correspond directly to elapsed time and each node is placed such that its position reflects its known or inferred date. Such trees are known as time trees, molecular clock phylogenies, or time stamped phylogenies. Furthermore, it became evident that substitution rates can vary between different parts of the tree and between sites along a sequence and several refinements of the original molecular clock tree have been developed. For a recent review of the evolution of methods for time tree estimation see (Kumar and Hedges, 2016).

In addition to questions regarding natural history, time trees are useful to study epidemiology and pathogen evolution (Gardy et al., 2015). Time trees of ‘measurably evolving’ pathogens can be used to date cross-species transmissions, introductions into geographic regions, and the time course of pathogen population sizes. In outbreak scenarios such as the recent Ebola virus or Zika virus outbreaks, rapid near real-time analysis of large numbers of viral genomes are necessary and can assist epidemiological analysis and containment efforts (Gardy et al., 2015).

BEAST is one of most sophisticated tools for time tree estimation (Drummond et al., 2012). BEAST samples many possible histories to evaluate posterior distributions of divergence times, evolutionary rates, and many other parameters. BEAST implements a large number of different phylogenetic and phylogeographic models. The sampling of trees, however, results in run-times of days to weeks for moderately large data sets of a few hundred sequences. On the other end of the spectrum are much simpler distance based tools that infer time scaled phylogenies orders of magnitudes faster (Britton et al., 2007; Tamura et al., 2012; To et al., 2016).

We developed a new tool called TreeTime that combines efficient heuristics with probabilistic sequence based inference. TreeTime infers maximum likelihood timetrees of a few thousand tips with within a few minutes. TreeTime was designed for applications in molecular epidemiology and analysis of rapidly evolving heterochronous viral sequences (Volz et al., 2013). It is already in use as an integral component of the real-time time outbreak tracking tool-kit nextstrain (Neher and Bedford, 2015). The main applications of TreeTime are ancestral state inference, evolutionary model inference, and time tree estimation. We discuss the core algorithms briefly below.

## Algorithms and implementation

TreeTime’s overarching strategy is to find an approximate maximum-likelihood configuration by iterative optimization of simpler subproblems similar in spirit to *sequential quadratic programming* or *expectation maximization*. Iteration is used on multiple levels, for example by iterating optimization of branch length, ancestral sequences, parameters of the relaxed clock, or coalescent models. Such an iterative procedure typically converges quickly when the branch lengths of the tree are short such that ancestral state inference has little ambiguity.

Maximum-likelihood assignments of sequence characters or node positions can be done such that the likelihood of a global state where all properties are specified jointly is maximal, such that the likelihood of local state after marginalization over all other unknown states is maximal. On a tree, both of these optimal assignments can be calculated in linear time (Felsenstein, 2004; Pupko et al., 2000) and TreeTime implements both marginal and joint ancestral reconstructions for ancestral states and node dates.

## Iterative branch length optimization

In general, optimizing the branch length of a tree is a complicated computational problem with 2*N* – 3 free parameters and a likelihood function that requires *O*(*N*) steps to evaluate. However, when branch length are ≪ 1, a joint optimization of branch length and ancestral sequences can be achieved by iteratively inferring branch length and ancestral sequences since corrections due to recurrent substitutions are neglibile. Given a tree topology and the branch length, the maximum likelihood ancestral sequences can be inferred in linear time (Felsenstein, 2004; Pupko et al., 2000). Likewise, maximum likelihood branch length given the parent and offspring sequences are easy to optimize. We use this iterative optimization scheme to rapidly optimize branch length and ancestral sequences. For more divergent sequences, however, integration over subleading states of internal nodes make a substantial contribution and the iterative optimization will underestimate the branch lengths. In this case, TreeTime can use branch length provided in the input tree.

## Maximum likelihood inference of divergence times

For a fixed tree topology and known ancestral sequences, the branch length corresponding to the maximum likelihood molecular clock phylogeny can be computed in linear time using dynamic programming or message passing techniques (Me´zard and Montanari, 2009). This approach is similar to Rambaut (2000), but the dynamic programming technique avoids computationally expensive numerical optimization of the branch lengths.

In analogy to maximum likelihood inference of ancestral sequence inference, the algorithm proceeds via a post-order tree traversal propagating the maximum likelihood assignments of subtrees towards the root, and a pre-order traversal selecting the optimal subtree given the constraints from above. Specifically, to calculate the joint maximum likelihood subtree, we calculate in post-order for each node *n*

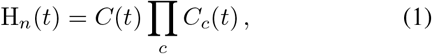

the likelihood that the node sits at position *t* given all constraints of its children. *C*(*t*) are contraints imposed on the node, while the product runs over all children *c* of node *n* and multiplies the integrated constraints of all subtending trees. These constraints are propagated along the branches of the tree via

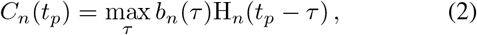

where *b*(*τ*) is the likelihood that the branch leading to the focal node *n* has length *τ*. Intuitively, *C_n_
*(*t_p_
*) specifies the distribution of the location *t_p_
* of the parent of node *n*, given the constraints from its tips and the substitutions that accumulated on the branch to the parent node.

During the post-order traversal, the branch length *τ* (*t_p_
*) maximizing Eq. 3 for *t_p_
* are tabulated and saved for the back-trace. Once the post-order traversal arrives at the root, the root is assigned the time argmax*t_C_
*(*t*) ∏_c_ *C_c_
*(*t*).

The post-order traversal is followed by a pre-order back-trace, during which the branch length of each internal node is assigned to the *τ* (*t_p_
*) conditional on the parental position *t_p_
*. To accelerate the optimization, TreeTime tabulates the branch length likelihood and the subtree likelihoods and creates linear interpolators of their logarithms.

The above algorithm assigns each node to the time that maximizes the joint likelihood of all branch lengths in analogy to the ancestral state reconstruction algorithm by Pupko et al.(2000). The marginally optimal time of each internal node, i.e., the time after integration over all other unconstrained nodes, can be determined in a similar manner by replacing the max in Eq. 3 by an integral over *τ*.

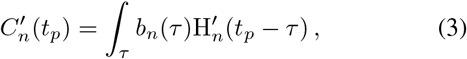

where H^′^ is the analog of H multiplying the *C^′^
* of all children. The marginal distribution of the time of a node is then given by

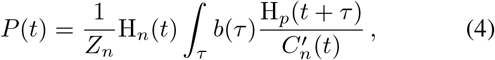

where *Z_n_
* is a normalization factor. Note that the contribution of node *n* to H_
*p*
_ is removed by dividing by it. TreeTime allows to compute joint and marginal maximum likelihood timings.

The result of the marginal reconstruction is a probability distribution of the node time, given the tree, ancestral sequence assignment, and the evolutionary model while the unknowns times of other nodes are traced out. From this distribution, arbitrary confidence intervals of node timing can be computed in a straight-forward manner.

The algorithm described above can be used for any continuous character on the tree. In Eq. 3, *b_n_
*(*τ*) can be replaced by and transmission function that depends either on the branch or properties of the child and parent node. We will use an analogous algorithm below to estimate parameters of relaxed molecular clock models.

## Efficient search for the optimal root

The fraction of variance in root-to-tip distance explained by a linear regression on sampling date is given by

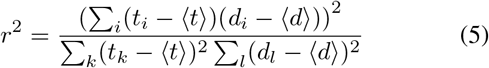

where the sum runs over all tips *i* of the tree and *t_i_
* and *d_i_
* are the sampling date and the distance from the root to node *i*, respectively. The angular brackets denote the sample average. The regression and *r*
^2^ depend on the choice of root via the *d_i_
*. In absence of an outgroup, the root is often chosen to maximize *r*
^2^ or minimize the squared residuals of a linear fit to the root-to-tip distance. Programs such as TempEst (Rambaut et al., 2016) and LSD (To et al., 2016) allow to search for the root that maximizes this correlation and TreeTime has implemented similar functionality.

This search for the optimal root can be achieved in linear time in the number of sequences *N* by first calculating for each internal node *n*

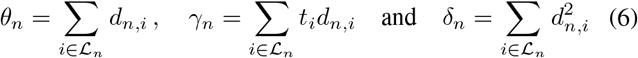

for each internal node *n*. Here, the sum runs over all tips *i ∈ L_n_
* of node *n* while *t_i_
* and *d_n,i_
* are the sampling date and the distance of tip *i* from node *n*, respectively. The quantities *θ_n_
*, *γ_n_
* and *δ_n_
* can be calculated recursively from the quantities of the children during one post-order traversal such that the entire calculation requires linear time, see appendix. Once those quantities are calculated, the corresponding quantities Θ_
*n*
_, Γ_
*n*
_, and ∆_
*n*
_ where the sum runs over all tips can be calculated in one pre-order traversal.

Hence *r*
^2^ for each internal node on the tree can be calculated in two tree traversals and the optimal choice determined in linear time. The mean squared residual can be calculated analogously.

In general, the optimal position of the root will not be an internal node, but a position between two nodes on a branch of the tree. Such optimal position on internal branches of the tree can be determined from the quantities calculated above by solving a quadratic equation without any numerical optimization. Details of the required algebra are given in the appendix.

## Resolving polytomies

Phylogenetic trees of many very similar sequences are often poorly resolved and contain multifurcating nodes also known as polytomies. Tree building software often randomly resolves these polytomies into a series of bifurcations. However, the order of bifurcations will often be inconsistent with the temporal structure of the tree resulting in poor approximations. To overcome this problem, TreeTime can prune all branches of length zero and resolve the resulting polytomies in a manner consistent with the sampling dates. For each pair of nodes, TreeTime calculates by how much the likelihood would increase when grouping this pair of nodes into a clade of size two. The polytomy is then resolved iteratively by always grouping pairs corresponding to the highest gain.

## Coalescent priors

The genealogical tree of individuals within a species depends on the size of the population, its geographic structure and fitness variation in the population (Kingman, 1982; Neher, 2013; Nordborg, 1997). In the simplest case of a panmictic population without fitness variation, the genealogies are described by a Kingman coalescent (Kingman, 1982), possibly with a population size that changes over time. Within the Kingman coalescent, any two lineages merge at random with a rate *λ*(*t*) that depends on the population size *N* (*t*) and the current number of lineages *k*(*t*).

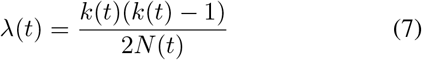

The rate at which a given lineage merges with any of the other lineages is *κ*(*t*) = (*k*(*t*) − 1)*/*2*T_c_
*(*t*). Here, the population size *N* (*t*) defines a time scale measured in units of generation time and we will more generally refer to this time scale by *T_c_
*(*t*) and measure it in units of the inverse clock rate.

The contribution of a branch between time points *t*
_0_ (child) and *t*
_1_ (parent) in the tree to the likelihood is then given by

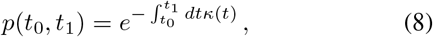

where a merger at time *t* contributes with rate *λ*(*t*)

TreeTime adds the contribution of each branch to the coalescent likelihood the branch likelihood object, which are then parameterized by the starting and end point of the branch, *b_n_
*(*t_n_, t_n_
* + *τ*). The total coalescent likelihood given a tree can be evaluated in one tree traversal such that *T_c_
* can be optimized efficiently. In addition to a constant *T_c_
*, TreeTime can model *T_c_
* as a piecewise linear function. Such piecewise functions are known as “skyline” (Strimmer and Pybus, 2001) and can be optimized by TreeTime as well.

## Relaxed clocks

Substitution rates can vary across the tree and models that assume constant clock rates may give inaccurate inference. Models that allow for clock-rate variation have been proposed (Drummond et al., 2006; Hasegawa et al., 1989; Yoder and Yang, 2000). These models typically regularize the clockrate through a prior and penalize rapid changes of the rate by coupling the rate along branches – known as autocorrelated or local molecular clock (Aris-Brosou et al., 2002; Thorne et al., 1998). TreeTime implements an autocorrelated molecular with a normal prior on variation in clock rates. The choice of the normal prior allows for an exact and linear time solution for the rate variation given the tree using the same forward/backward trace algorithm used for the inference of internal nodes.

## Inference of time reversible substitution models

Large phylogenies typically contain 100s of substitutions and thus provide enough information to infer substitution models from the data. TreeTime implements an iterative algorithm to infer general time reversible substitution models (Felsenstein, 2004) parameterized by equilibrium state frequencies *π_i_
* and a symmetric substitution matrix *W_ij_
*. The instantaneous rate from state *j i* is *Q_ij_
* = *π_i_W_ij_
*. The model is inferred by first counting the time spend in different states across the tree *T_i_
* and the number of substitutions between *n_ij_
* in a joint maximum likelihood assignment using a simple substitution model. Then, *π* and *W* are determined by iterating

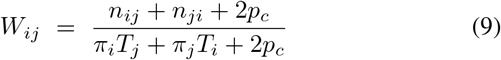

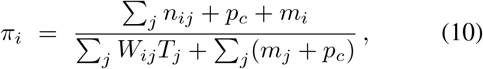

where *p_c_
* is a small pseudo-count driving the estimate towards a flat Jukes-Cantor model in absence of data, and *m_i_
* are the number times state *i* is observed in the sequence of the root. In each iteration, the *π* is normalized to one, the diagonal of *W_ij_
* is set to 
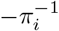
Σ_i≠j_
*W_ij_π_j_ W_ij_
* is rescaled such that the total expected substitution rate -∑*π_i_W_ii_π_i_
* equals one. The rescaling of *π* and *W_ij_
* can be absorbed into an overall rate *µ*. This algorithm typically converges in a few iterations.

## Implementation

TreeTime is implemented in python and uses the packages numpy and scipy for optimization, linear algebra, and interpolation (Jones et al., 2001–2017; van der Walt et al., 2011). Computationally costly operations are cast into array operations executed by numpy when ever possible. Tree-Time is organized as a hierarchy of classes. TreeAnc performs maximum likelihood inference of ancestral sequences, ClockTree infers a time scaled phylogeny given a tree topology, TreeTime adds an additional layer of functionality including rerooting, polytomy resolution, coalescent models, and relaxed clocks. The substitution model is implemented as a general time reversible model in the class GTR.

This structure allows TreeTime to be used in a modular fashion in python based phylogenetic analysis pipelines. In addition, scripts can be called from the command line to perform standard tasks such as ancestral character inference, re-rooting of trees, and time tree estimation.

## Availability

TreeTime is published under an MIT license and available at github.com/neherlab/treetime. The complete data sets used to validate TreeTime are available at https://io.scicore.unibas.ch/index.php/s/raTptOICG9MsDIO.

## Validation and performance

To assess the accuracy of date reconstructions of treetime and to compare its performance to existing tools such as BEAST and LST (Drummond et al., 2012; To et al., 2016), we generated toy data using the FFPopSim forward simulation library (Zanini and Neher, 2012). We simulated population of size *N* = 100 and used a range evolutionary rates *µ = 10 ^-5^,…,0.002* resulting in expected genetic diversity from 0.001 to 0.2. Sequences were sampled every 10, 20, or 50 generations. The length of the simulated sequences was *L* = 1000.

Fig. 1 shows the error in the estimates of the clock rate for TreeTime, LSD, and BEAST as a function of the evolutionary rate. TreeTime and LSD estimates of the clock rate are very for small rates but tend to underestimate the rates at when diversity exceeds a few percent. This is expected, as maximum likelihood inference underestimates branch lengths. BEAST produces accurate estimates across the entire range of diversities. By sampling trees, BEAST does not suffer from the atypical maximum likelihood assignments.

**FIG. 1.**
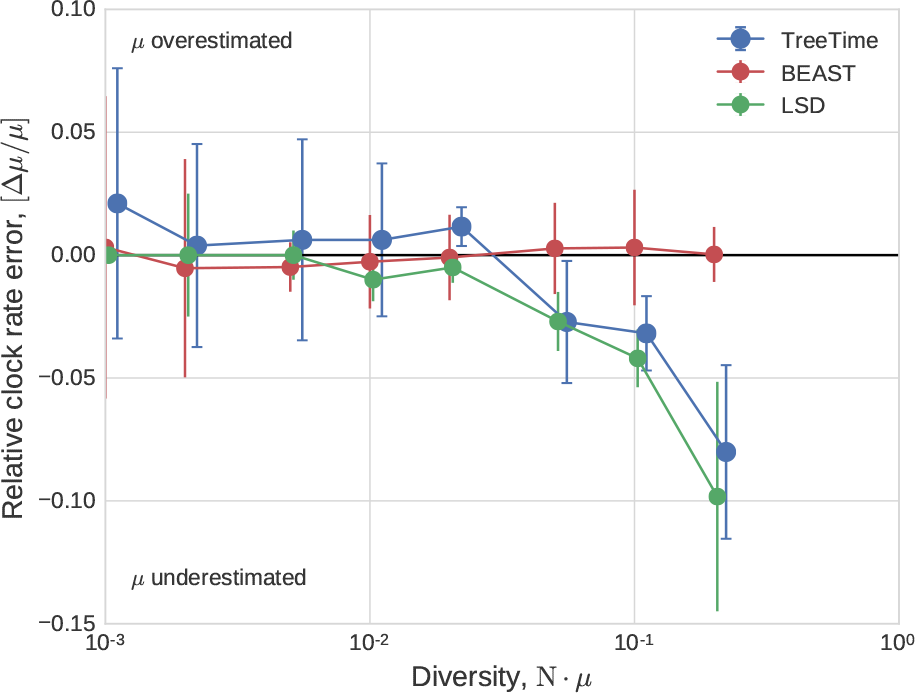
Estimation of the evolutionary rate from simulated data by TreeTime, LSD, and BEAST. TreeTime and LSD (after tree reconstruction using FastTree) underestimate the rate when branch length are long. The error bars denote *±* one standard deviation.

**FIG. 2.**
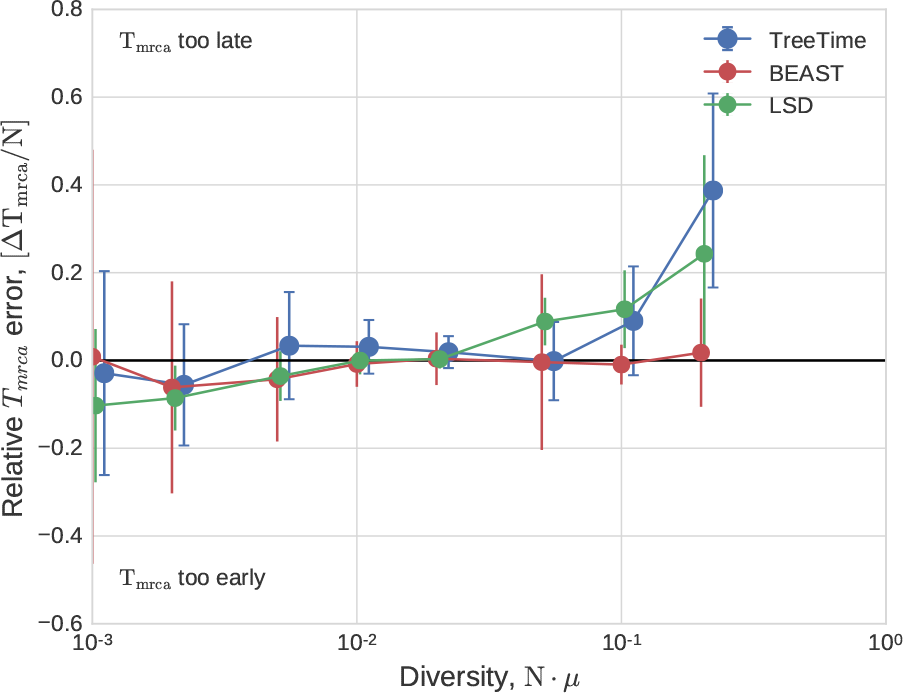
Estimation of TMRCA from simulated data by TreeTime, LSD, and BEAST. All three programs estimate the time of the MRCA with 10% accuracy, except for the very long branches when TreeTime tends to overestimate the age of the root. Error bars show one standard deviation.

In a similar manner, TreeTime, LSD, and BEAST estimate the time of the most recent common ancestor to within 10% accuracy (relative to the coalescence time) across the range of simulated data.

We also ran TreeTime on simulated data provided by To et al.(2016) and compared it to the results reported by To et al. (2016) for LSD, BEAST and a number of other methods. Fig. 3 compares the accuracy of *T_M RCA_
* and clock rate estimates, showing that TreeTime achieves similar or better accuracy than other methods.

**FIG. 3.**
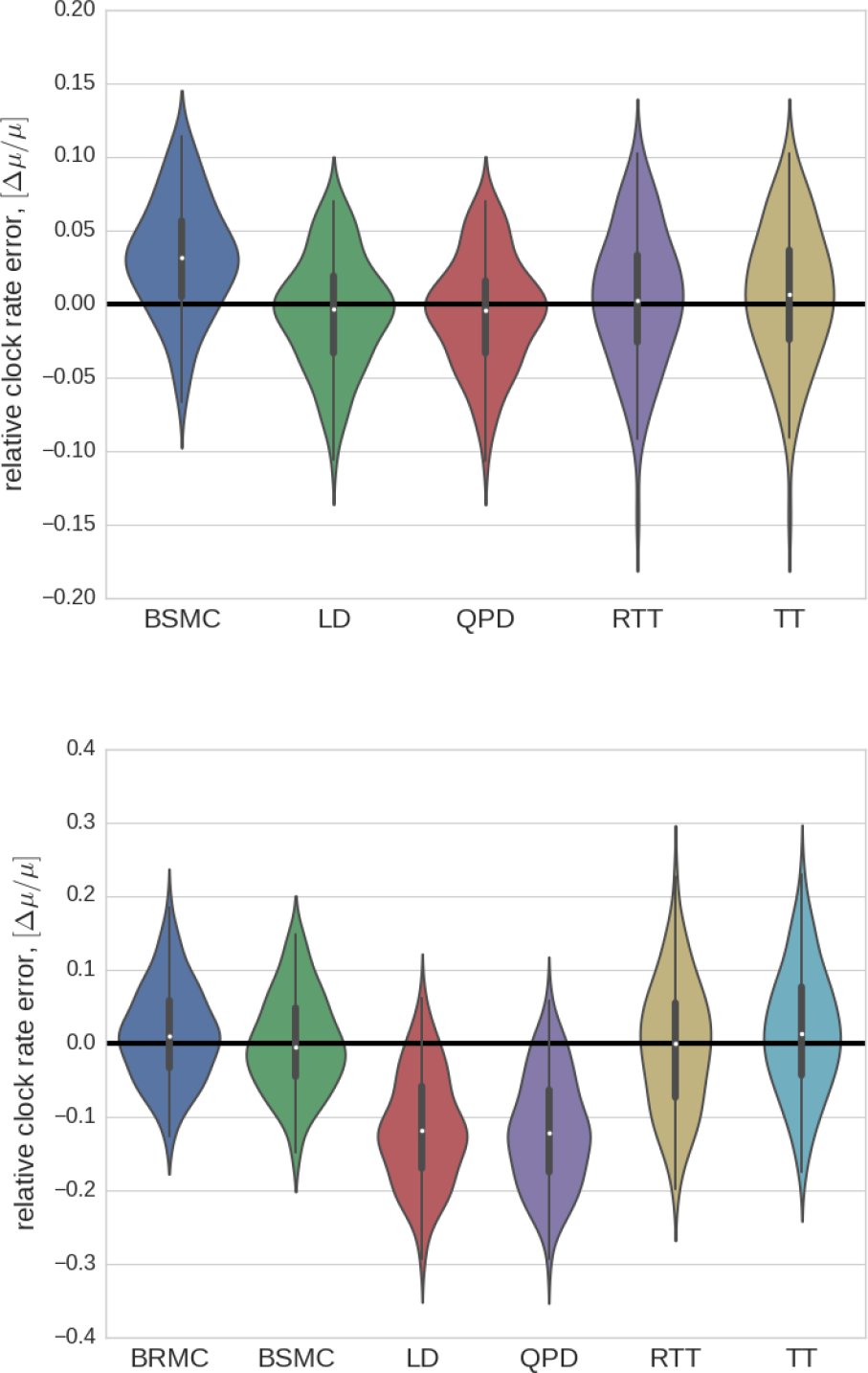
**LSD test data.** TreeTime has comparable or better accuracy as BEAST (BSMC), LSD (LD, QPD), or root-to-tip regression (RTT) when run on simulated data provided by (To et al., 2016). Both panel use the tree set 750_3_25, the top and bottom panel show runs on alignments generated with a strict and relaxed molecular clock, respectively.

*a. Coalescent model inference:* Population bottlenecks, selective sweeps, or population structure, affect the rate of coalescence in an often time variable way. BEAST can infer a history of effective population size (inverse coalescent rate) from a tree – often known as skyline. TreeTime can do a similar inference by maximizing the coalescence likelihood with respect to the pivots of a piecewise line approximation of the coalescence history *T_c_
*(*t*). To test the power and accuracy of this inference, we simulated sinusoidal population size histories of different amplitude and period, uniformly sampled sequences through time, and used these data to estimate the coalescent rate history. Comparisons of true and estimated histories are shown in Fig. 4.

**FIG. 4.**
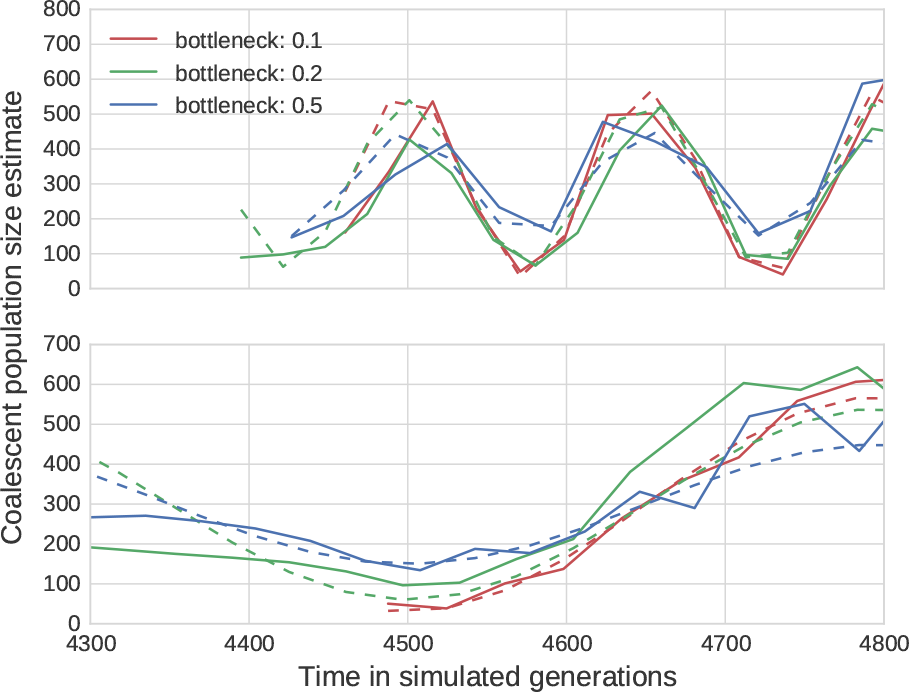
Reconstruction of fluctuating population sizes by TreeTime. The graph shows simulated population size trajectories (dashed lines) and the inference by TreeTime as solid lines of the same color. Different lines vary in the bottleneck sizes of 10%(red), 20%(green) and 50%(blue) of the average population size. The top panel shows data for fluctuations with period 0.5*N*, the bottom panel 2*N*. The average population size is *N* = 300.

## Influenza phylogenies

The dense sampling of influenza A virus sequences over many decades makes this virus an ideal test case to evaluate the sensitivity of time tree estimation to sampling depth. We estimated the clock rate and the time of the most recent common ancestor of influenza A H3N2 HA sequences sampled from 2011 to 2013 for sets of sequences varying from 30 to 3000, see Fig. 5. TreeTime estimates are stable across this range, while estimates by LSD tend to drift with lower rates and older MRCAs for larger samples. Estimates by BEAST are generally concordant with TreeTime.

**FIG. 5.**
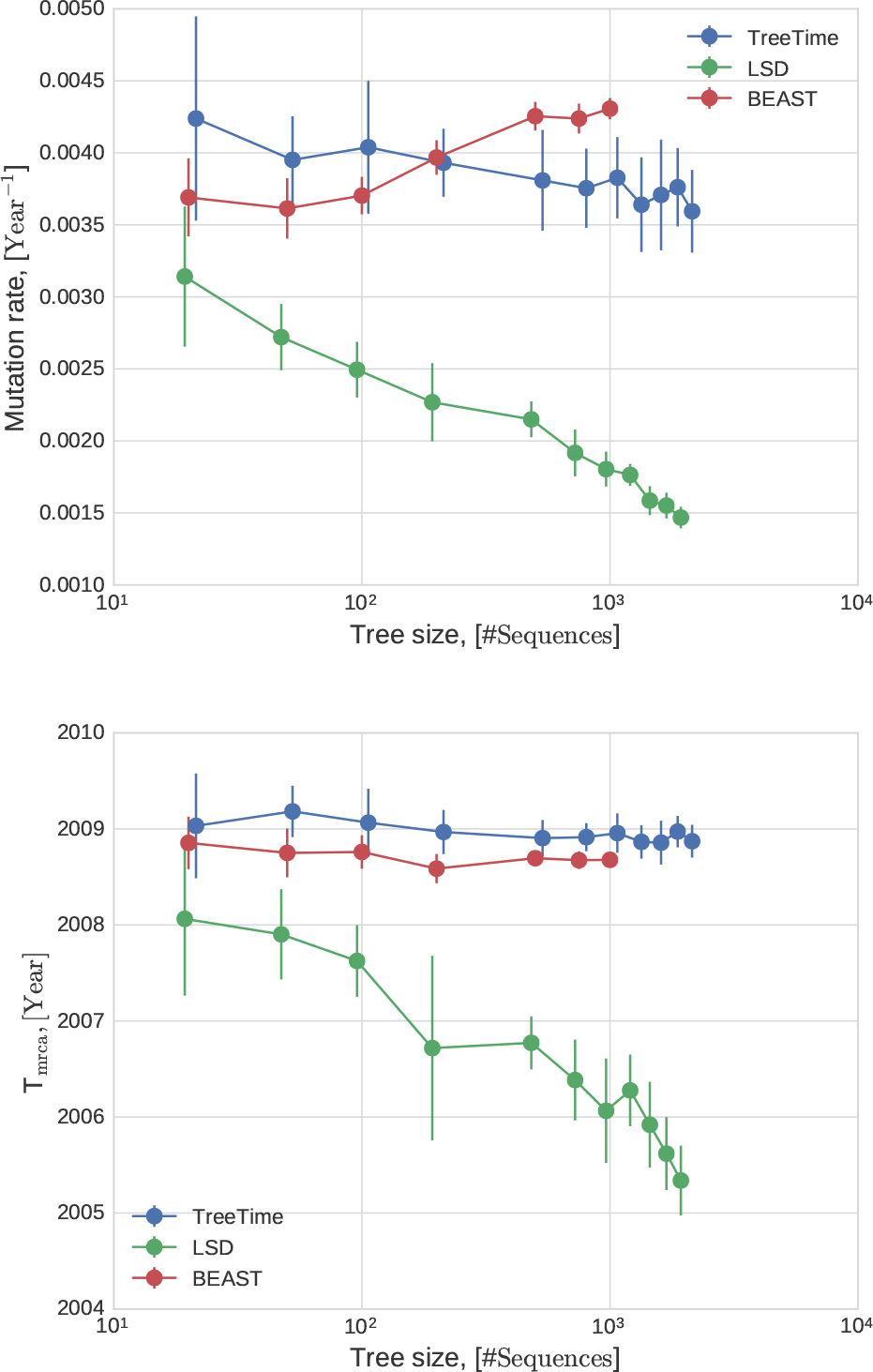
Variation of the estimate of the rate of evolution of H3N2 and the for different sensities of sampling

Next, we tested how accurately TreeTime infers dates of tips when only a fraction of tips have dates assigned. Every tip in TreeTime can either be assigned a precise date, an interval within which the date is assumed to be uniformly distributed, or no constraint at all. TreeTime will then determine the probability distribution of the date of the node based on the distribution of the ancestor and the substitutions that occurred since the ancestor. We tested the accuracy at which missing dates can be inferred in an influenza phylogeny by erasing date information of a fraction (5% to 95%) of all nodes, see Fig. 6.

**FIG. 6.**
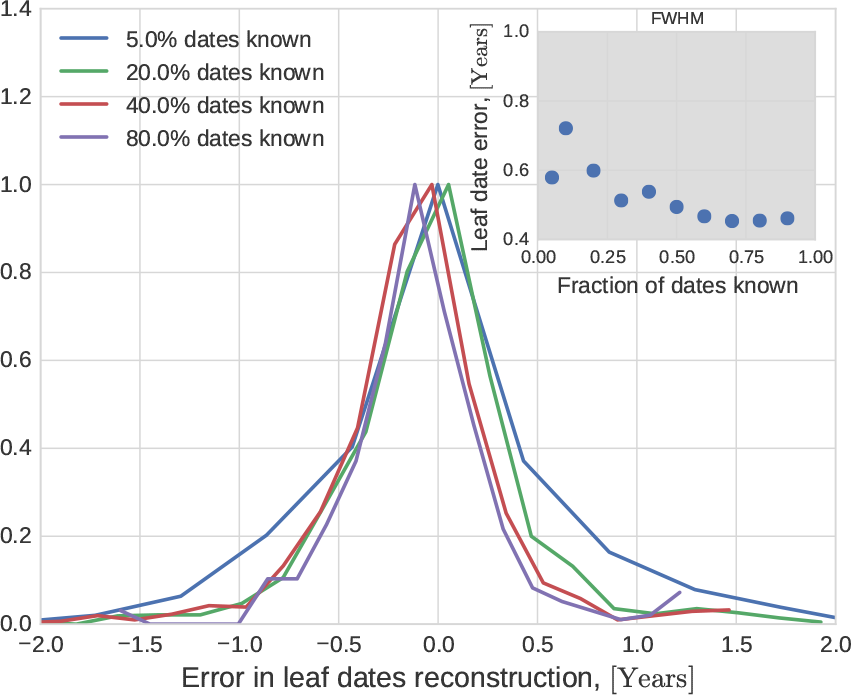
Tip dating and sensitivity to missing information. A) The inter-quartile range of the error of estimated tip dates decreases from 0.7 years to 0.5 years as the fraction of known dates increases from 5% to 90% (see inset).

In summary, on data sets with short branches but fairly unambiguous topologies, timetrees inferred by TreeTime have similar accuracy to those inferred by BEAST but results are obtained in a fraction of the time.

## Analysis of the 2014-2015 Ebola Virus outbreak

In 2014, West Africa experienced the largest known outbreak of Ebola Virus (EBOV) in humans. The genomic epidemiology has been studied intensively by multiple groups (Dudas et al., 2017). Here, we reanalyzed a subset of 350 EBOV sequences sampled throughout the outbreak from 2014-2016. Due to the dense sampling, the maximum likelihood phylogeny has many unresolved nodes and TreeTime was used to resolve polytomies using temporal information. After automatic rooting and GTR model inference, TreeTime produced the timetree shown in Fig. 7. The GTR model inferred from the tree was

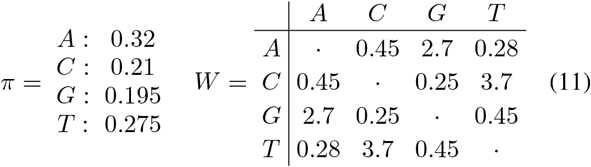

**FIG. 7.**
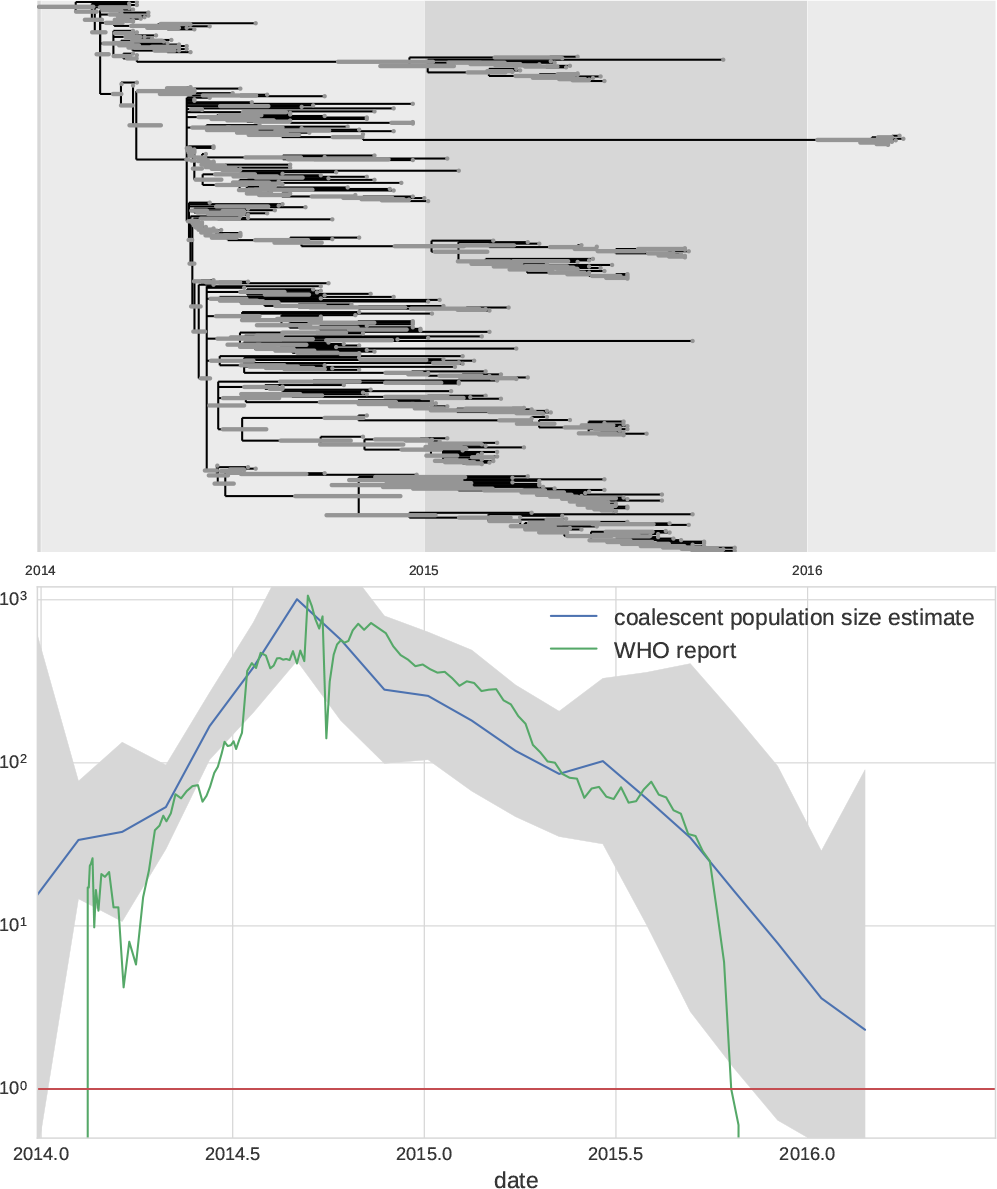
**EBOV phylodynamic analysis.** The top panel shows a molecular clock phylogeny of EBOV sequences obtained over from 2014-2016 in West Africa. The lower panel shows the estimate of the coalescent population size along with its confidence intervals. The estimate suggest an exponential increase until late 2014 followed by a gradual decrease leading to almost complete eradication by 2016. Ebola case counts, as reported by the WHO (2016) agree quantitatively with the estimate.

TreeTime ran 4min on a regular laptop to complete this analysis. In addition to inferring a time tree, TreeTime estimated the time course of the coalescent population size shown im the lower panel of Fig. 7. The estimated population size closely mirrors the case counts reported by the WHO throughout this period.

## Discussion

TreeTime was developed with large heterochronous viral sequence alignments in mind and we have used TreeTime as the core component of the nextstrain real-time phylogenetics pipeline (Neher and Bedford, 2015). TreeTime tries to strike a useful compromise between inflexible but fast heuristics and computationally expensive Bayesian approaches that require extensive sampling of treespace. The overarching algorithmic strategy is iterative optimization of efficiently solvable sub-problems to arrive at a consistent approximation of the global optimum. While this strategy is approximate and often assumes short branch length, it converges fast for many applications and trees with thousands of tips can be analyzed in a few minutes.

Rapid and efficient analysis methods are of increasing importance as data sets are increasing in size. During the recent outbreaks of EBOV and Zika virus, hundreds of sequences were generated and need to be analyzed in near real time to inform containment efforts. Similarly, the GISRS network for surveillance of seasonal influenza virus sequences hundreds of viral genomes per month. Timely analysis of these data with Bayesian methods such as BEAST is unfeasible. TreeTime addresses this need.

Compared to other methods recently developed for rapid estimations of timetrees (Britton et al., 2007; Tamura et al., 2012; To et al., 2016), treetime uses probabilistic models of evolution, allows inference of ancestral characters, and coalescent models. In TreeTime, every node of the tree can be given a strict or probabilistic date constraint. This higher model complexity results in longer run times, but the scaling of run times remains linear in the size of the data set and alignments with thousands of sequences can be analyzed routinely. The timetree inference and dating are typically faster than the estimation of the tree topology.

TreeTime can be used in a number of different ways. The core TreeTime algorithms and classes can be used larger phylogenetic analysis in python scripts. This is the most flexible way to use TreeTime and all the different analysis steps can be combined in custom ways with user specified parameters. In addition, we provide command-line scripts for typical recurring tasks such as ancestral state reconstruction, rerooting to maximize temporal order, and time tree inference. We also implemented a web-server that allows exploration and analysis of heterochronous alignments in the browser without the need to use the command-line.

TreeTime was tested predominantly on mildly diverged sequences from viruses. The iterative optimization procedures are not expected to be accurate for trees were the many sites are saturated. In such scenarios with extensive uncertainty of ancestral states and tree topology, convergence of the iterative steps can not be guaranteed. While in many cases TreeTime might still give approximate branch length and ancestral assignments and timetree estimates, these need to be checked for plausibility. In general global optimization and sampling of the posterior can not be avoided.

## Appendix: Finding the optimal root in linear time

To calculate the correlation between the root-to-tip distances and tip dates via equation 5, one first needs to calculate the means and (co)variances of tip dates and root-to-tip distances. For a tree with *N* tips, this requires (*N*) operations and calculating it for all internal nodes would therefore require *
*O*
*(*N* ^2^) operations. The same covariances are needed to calculate the regression parameters and the residuals. However, its is possible to calculate the quantities for every node at once, reducing the total number of operations to (*ON*).

The speed-up is possible through recursively calculating sums and averages on the tree. We denote the set of tips that descend of node *n* by *L_n_
*. We will need the number tips *M_n_
* = | *L_n_
* | the sum of their sampling times *Τ_n_
* =Σ_
*i∈Ln*
_ *t*
_
*i*
_ the sum of their distances *d_n,i_
* from node *n θ_n_
* =Σ_
*i∈Ln*
_ d_
*n,i*
_, and the analogous higher order quantities *γn* =Σ_
*i∈Ln*
_ t_i_ d_n,i_ and 
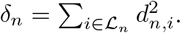

First, assign M_n_ = 1, τ_n_ = t_n_, Τ_n_ = 0, n = 0 and δ_n_ = 0 for all tips of the tree. Then, in one post-order transversal over internal nodes, we can calculate these quantities by summing the following expressions over the children C_n_ of node n.

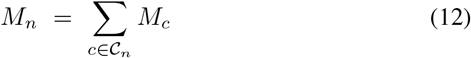

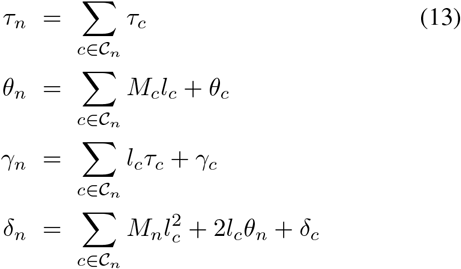

To calculate the covariances at a particular node *n*, we need to sum over all terminal nodes rather than only tips that descend from the node. We denote the corresponding quantities by capital letters. The sums of the sampling dates and their squares are of course straightforward to evalulate, the remaining quantities that depend on the choice of the focal node *n* can be calculated in one pre-order transversal. Let *p* denote the parent node of node *n*

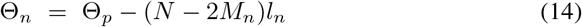

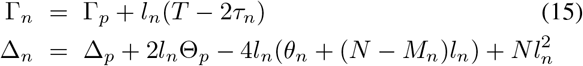

Note, that the order in which these calculations are performed matters. The first line, calculating Θ_
*n*
_ adjusts the parent value Θ_
*p*
_ for the fact that the branch leading to node *n* is transversed by *N - M_n_
* path instead of *M_n_
* if the root is shifted from *p* to *n*. Similarly, Γ_
*n*
_ is calculated from Γ_
*p*
_ by adjusting with the difference of sum of times of subtending and complementary nodes. The corresponding expression for the sum of squared root-to-tip distances is slightly more complicated but still follows from elementary algebra.

With these quantities at hand, the regression, residuals, and *r*
^2^ can be straightforwardly calculated from the means and covariances given by

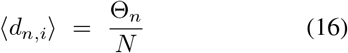

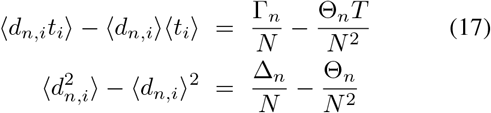

In genernal, the optimal root is not going to coincide with a preexisting node but will be placed somewhere along a branch. When placing the root at a position ∊ *∈* [0, 1] along the branch, the corresponding Θ_
*n*
_(*∈*), Γ_
*n*
_(*∈*), ∆_
*n*
_(*∈*) are obtained by substitution *Elc* for *lc* in equation 14. The fraction of variance explained by a root-to-tip regression with a root placed at position ∈ on a branch then has the generic form

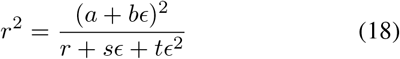

where the coefficients can be obtained by sustituting the expressions for Θ_
*n*
_(*∈*), Γ_
*n*
_(*∈*), and ∆_
*n*
_(*∈*). The condition for a maximum 
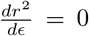
 results in an quadratic equation for ∈.Hence, the optimal position of the root can be calculated with a number of operations that increases linearly in the size of the tree.

The slope of the root-to-tip regression or clock rate in then simply 
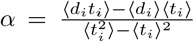
, where *d_i_
* are evaluated with respect to the optimal root.

